# Isolation and characterization of *Ovine gammaherpesvirus type 2* from an outbreak of Malignant Catarrhal Fever in *Artiodactyla* and horses in Mexico

**DOI:** 10.1101/2022.12.18.520964

**Authors:** Tania Lucia Madrigal-Valencia, Manuel Saavedra-Montañez, Armando Pérez-Torres, Jesús Hernández, Joaquim Segalés, Yesmín Domínguez Hernández, Irma Eugenia Candanosa-Aranda, Alfredo Pérez-Guiot, Humberto Ramírez Mendoza

**Affiliations:** Departamento de Microbiología e Inmunología, Facultad de Medicina Veterinaria y Zootecnia, UNAM, Mexico City, Mexico; Departamento de Biología Celular y Tisular. Facultad de Medicina, Universidad Nacional Autónoma de México (UNAM), Mexico City, Mexico; Laboratorio de Inmunología, Centro de Investigación en Alimentación y Desarrollo, A.C. (CIAD), Hermosillo, Sonora, Mexico; Unitat Mixta d’Investigació IRTA-UAB en Sanitat Animal, Centre de Recerca en Sanitat Animal (CReSA), Campus de la Universitat Autònoma de Barcelona (UAB), 08193 Bellaterra, Barcelona, Catalonia, España; Department de Sanitat i Anatomia Animals, Facultat de Veterinària, Campus de la Universitat Autònoma de Barcelona (UAB), 08193 Bellaterra, Barcelona, Catalonia, España; Centro de Enseñanza, investigación y Extensión en Producción Animal en Altiplano (CEIEPAA), Facultad de Medicina Veterinaria y Zootecnia (FMVZ), Universidad Nacional Autónoma de México (UNAM), Tequisquiapan, Queretaro, Mexico

**Keywords:** Horses, malignant catarrhal fever, Mexico, OvHV-2, primary cell cultures

## Abstract

Ovine gammaherpesvirus 2 (OvHV-2), a member of the Macavirus genus, causes sheep-associated malignant catarrhal fever (SA-MCF), a fatal lymphoproliferative disease that affects a wide variety of ungulates in addition to horses.This study described an outbreak of SA-MCF that occurred in Mexico and the identification of the OvHV-2 virus through viral isolation and different laboratory techniques such as immunofluorescence (IF), immunoperoxidase (IP), immunohistochemistry (IHC), end point PCR and partial sequencing of the ORF75 gene. The animals involved in this outbreak showed head and eye clinical signs and lesions. Based on the clinical-pathological outcome, buffy coats were taken, and virus isolation was attempted on primary cell cultures of the rabbit testicle. Small clusters of refractile cytomegalic cells characterized the cytopathic effect between 48 and 72 hours postinfection. In addition, inclusion bodies were identified, and cytoplasmic immunoreactivity was observed in the infected cells. The sequences obtained were aligned with OvHV-2 sequences reported in GenBank and revealed a nucleotide identity higher than 98%. The results indicate that the outbreak was caused by OvHV-2 and the horses are susceptible to SA-MCF.

## 1. Introduction

Malignant catarrhal fever (MCF) is a severe lymphoproliferative disease syndrome in susceptible ungulate species of the order Artiodactyla. MCF may be caused by any of the gammaherpesviruses that comprise the *Macavirus* classification group. The two most well-studied are ovine herpesvirus-2 (OvHV-2) and alcelaphine herpesvirus-1 (AIHV-1), which are maintained asymptomatically in sheep and wildebeest reservoir populations, respectively (1). OvHV-2 is responsible for sheep-associated MCF (SA-MCF) disease in susceptible hosts such as buffalo (2), cattle (3), and deer (4) and rarely in pigs (3) and foals (5). In OvHV-2, infection of nonadapted host species suggests latent infection and abortive lytic viral replication (6).

Typical SA-MCF, often referred to as the head-and-eye form, is the most common presentation in cattle (1). Animals develop pyrexia, anorexia, bilateral corneal opacity, nasal and ocular discharge, ulceration on the mucosa, and neurological manifestations in terminal stages (7). The most common gross pathological changes in the SA-MCF affected animals are erosions of the tracheal and bronchial mucosa, erythema of the turbinate mucosa, congestion and edema of the lungs, and focal white lesions in the kidney (7). Histopathologically, the hallmark of SA-MCF is lymphoproliferative inflammation with vasculitis involving medium-caliber arteries and veins, which is readily detected in most organs of cattle dying of acute SA-MCF (8).

For the in vitro isolation of MCF-causing viruses, primary cell cultures of bovine thyroid (BTh), bovine kidney (BK), bovine embryonic kidney (BEK) and calf testis (CT) cells are used (9–11).

The diagnosis of MCF is made when typical clinical signs and lesions are observed, combined with a recent history of exposure to a reservoir host (12). For the detection of antibodies against MCF virus (MCFV), several serological assays have been developed, such as enzyme-linked immunosorbent assay (ELISA), competitive inhibition ELISA (cELISAv), (13,14), immunofluorescence (IF), immunoperoxidase (IP), and the complement-fixation test (15,16). However, the IF, IP and complement-fixation tests use AlHV-1 as an antigen. PCR is a sensitive and specific test that can support the diagnosis of SA-MCF(17). Buffy coats and lymphoid tissues are ideal sources to isolate the OvHV-2 virus and to perform PCR tests (18).

SA-MCF cases have been documented in North America, and the losses due to this disease forced bison producers to leave the market (19). In 2003, more than 800 heads of bison were lost, representing financial losses of approximately one million dollars (20).

Regarding the history of MCF in Mexico, in 1969, the first case of SA-MCF was recognized in Mexico City in a 6-year-old Holstein cow, which presented a head and ocular clinical manifestation, in addition to perivascular lymphoid infiltrates in various organs. However, the diagnosis was only based on a clinical-pathological study (21). In this context, an outbreak of SA-MCF occurred in the municipality of Tequisquiapan in the state of Queretaro, which affected many animals with a head and ocular clinical presentation.

Only one study has been carried out with OvHV-2 in Mexico, since it is considered an exotic disease and the economic and epidemiological impact, as well as the distribution and molecular characterization of the OvHV-2 virus in this country are unknown. Therefore, the objective of this study was to isolate and identify the OvHV-2 virus that caused the SA-MCF outbreak in Mexico.

## 2. Material and Methods

### 2.1 Samples

Sampling was performed at the Centro de Enseñanza, Investigación y Extensión en Producción Animal en Altiplano (CEIEPAA), located in Tequisquiapan, state of Queretaro, Mexico. This farm is based on a semi-intensive production system with multiple species (deer, dairy cattle, feedlot cattle, sheep, goats, and horses) (Figure 1).

**Figure 1.**
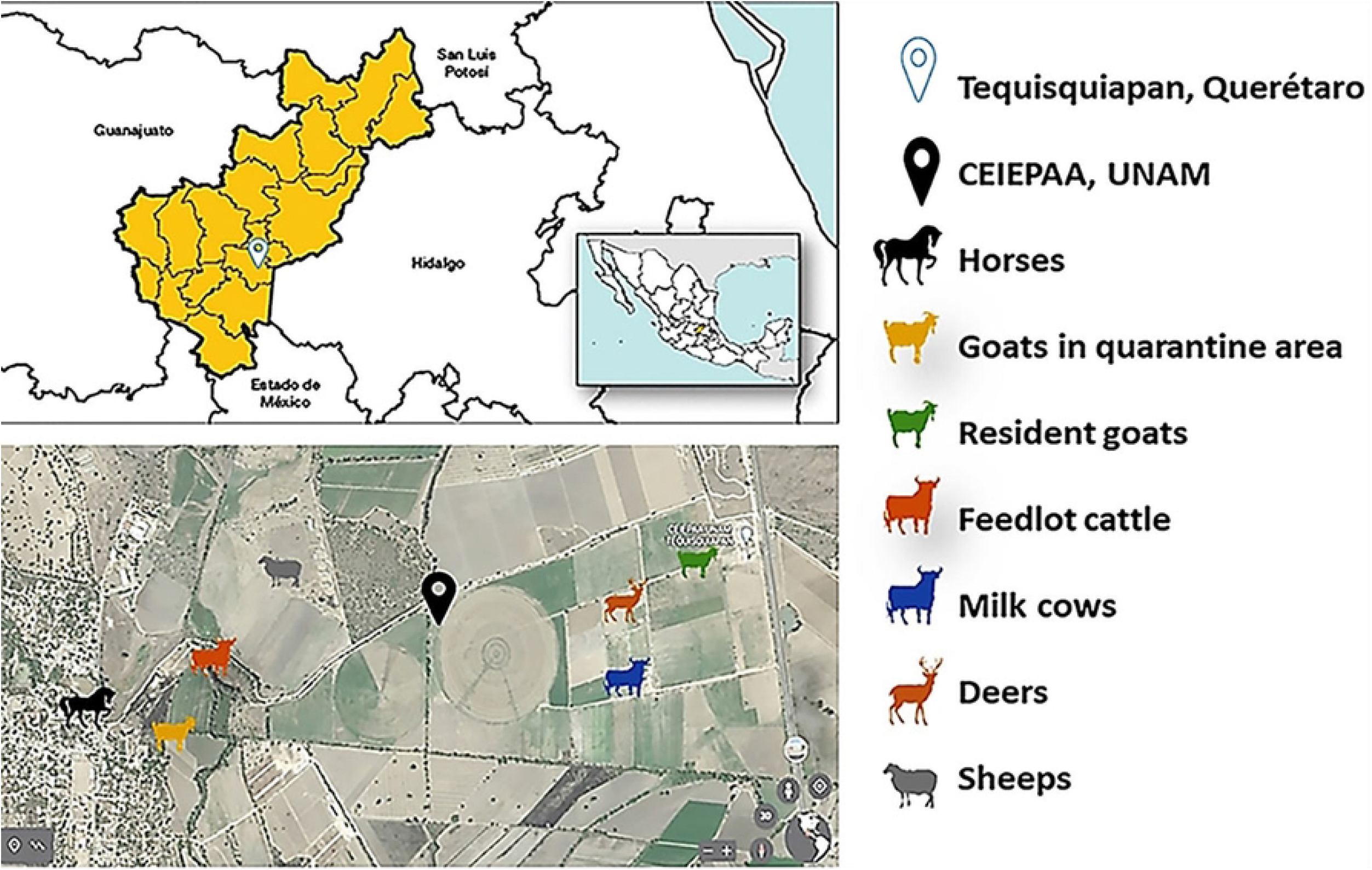
Distribution of animals in the CEIEPAA farm, UNAM. On the upper left, the geographical location of Tequisquiapan, Queretaro (Mexico), is indicated. The species distribution in the area is depicted on the left inferior section.

The sampled artiodactyls had vesicular lesions in the oral mucosa, hypersalivation and corneal opacity, except for the goats and sheep, which did not present any clinical signs. In addition, these animals were PCR or RT–PCR negative for foot and mouth disease (FMD), infectious bovine rhinotracheitis (IBR), vesicular stomatitis serotypes Indiana and New Jersey (VS), parapoxvirus (PR), bluetongue (BT) and contagious ecthyma (CE) viruses. The sampled horses displayed the same clinical signs and lesions and were negative for vesicular stomatitis virus by RT–PCR and isolation (Table 1).

**Table 1.** Affected species, positive by PCR, percentage of positive animals.

| Affected species | *Previous Diagnoses/Results (PCR) | **Clinical signs | Total population/sick/dead | Sampled animals, PCR-positive and percentage of positive |
| --- | --- | --- | --- | --- |
| Horses | VS/(-) | MUMV,UT,LF<br>C,WL | 39/14/0 | 39 [9] (23%) |
| Feedlot cattle | IBR,VS,PR,FMD,<br>BT/(-) | MUMV,HS,LF<br>C,WL | 87/7/0 | 30 [11] (36%) |
| **Dairy cows** | IBR,VS,PR,FMD,<br>BT/(-) | MUMV,HS,LF<br>C,WL | 163/23/0 | 32 [5] (15%) |
| **Resident goats** | IBR,VS,PR,FMD,<br>BT/(-) | - | 297/0/0 | 32 [19] (59%) |
| **Goats in**<br>**quarantine area** | IBR,VS,PR,FMD,<br>BT,CE/(-) | - | 97/0/0 | 32 [8] (25%) |
| **Sheep** | IBR,VS,PR,FMD,<br>BT,CE/(-) | - | 222/0/0 | 30 [26] (86%) |
| **Deer** | IBR,VS,PR,FMD,<br>BT,CE/(-) | MUMV,HS,CO<br>,LFC,WL | 82/31/14 | 18 [8] (44%) |
| **Total** |  |  | **987/117/14** | **213 [86] (40.3%)** |

Results of the nested end-point PCR for detecting the OvHV-2 virus in buffy coat samples of different species.
[ ] = positive to PCR, ( ) = percentage of animals positive to PCR.
\*Infectious Bovine Rhinotracheitis (IBR), Vesicular Stomatitis serotypes Indiana and New Jersey (VS), Parapoxvirus (PR), Foot and Mouth Disease (FMD), Bluetongue (BT), Contagious Ecthyma (CE).
\*\*mucogingival ulcers in the mouth vestibule (MUMV), ulcers in the tongue (UT), hypersalivation (HS), corneal opacity (CO), lower food consumption (LFC), and weight loss (WL).

Deer that died during the outbreak in CEIEPAA have undergone routine evaluations shortly after death. Cornea, lymph node, tongue, rumen, liver, and lung tissue samples were fixed by immersion in buffered formalin solution at 10%, routinely processed for histological evaluation and visualized with Hematoxylin and eosin (H&E).

Approximately 30 blood samples were obtained from each animal species, which were used to separate buffy coats according to the methodology described by English & Andersen (1974) to attempt viral isolation. Samples were identified and stored at −70°C until processing (22).

### 2.2 Cell cultures

Primary cell cultures of rabbit testicles were used to isolate and identify OvHV-2. Two-month-old male rabbits were selected and humanely euthanized per the Institutional Subcommittee for the Care and Use of Experimental Animals of the UNAM (SICUAE.DC-2020/3-5). For the processing of the testes, all external connective tissue layers were first removed and cut into small pieces, and trypsin-EDTA was added at a 1:5 proportion. Then, fragments were incubated at 37°C for 30 min, and finally, the supernatant was centrifuged at 800 × *g* for 10 min. The cells were suspended in Dulbecco’s modified Eagle’s medium (DMEM) supplemented with 10% bovine fetal serum and incubated at 37°C in a 5% CO_2_ atmosphere.

All positive buffy coats obtained from the above-described animals with clinical signs of SA-MCF and identified by PCR were used for inoculating cell cultures. These buffy coats were pooled by species and processed separately. The cell cultures were inoculated with 200 μL of each mixture and propagated in DMEM supplemented with 2% bovine fetal serum and incubated at 37°C for 96 h. At least four blind passages were performed, and after the cytopathic changes of the monolayers, the isolated agents were identified for OvHV-2 virus DNA by nested PCR. The viral titer was calculated with the Reed and Muench method (23).

To study the inclusion bodies, a 24-well plate with an 80% confluence cell culture was used. Each well was infected with 200 μl of each blind passage until completing the four blind passages of the six isolates. Subsequently, the plate was incubated for 24 hours at 37°C with 5% CO2. Afterwards, the supernatant was removed and fixed with 4% paraformaldehyde. Finally, hematoxylin was added and it was allowed to incubate for 1 minute, washed with distilled water and eosin dye was added, and it was allowed to incubate for 30 seconds. Finally, glycerol was added and each well was covered with a coverslip and observed under a microscope. As a negative control, a 4-well plate was left with uninfected cells, which were stained with H&E.

### 2.3 Hyperimmune serum

To obtain a hyperimmune serum against OvHV-2, three rabbits were inoculated intramuscularly with 1 mL of the viral isolation from horses (10^4.19^ TCID_50_ / mL) every week for 5 weeks; after cardiac puncture from three rabbits, serum was obtained, pooled and finally stored at −20°C.

### 2.4 Ethics Statements

All of the experimental procedures involving animals were conducted in accordance with the Institutional Subcommittee for the Care and Use of Experimental Animals of the UNAM (SICUAE.DC-2020/3-5).

### 2.5 Immunoperoxidase (IP) and indirect immunofluorescence (IF) techniques

Primary cell cultures with 80% cell confluency were used for the IF and IP techniques. These primary cell cultures of rabbit testis cells were inoculated with 50 μL of the third passage from each of the six isolates separately, with a minimum titer of 10^3^ to 10^4.19^ TCID _50_%_/_mL in 96-well plates. Then, they were incubated at 37°C for 72 h and fixed with 4% paraformaldehyde.

The plates for IP were blocked with 3% hydrogen peroxide, and the plates for IF were blocked with 2.5% bovine serum albumin, adding the polyclonal rabbit antibody at a dilution of 1:5. The plates were washed with phosphate-buffered saline (PBS), the conjugate (peroxidized protein A) was added for IP at a 1:16 dilution, and fluorescein isothiocyanate (FITC) was added at a 1:20 dilution for IF. Plates were incubated at 37°C for 1 hour, and the plate was supplemented with the diaminobenzidine substrate (15). After 5 min, the reaction was stopped with distilled water. The antigen-antibody reaction was determined as an insoluble brown coloration in the IP plates and a green-apple-colored fluorescent emission at 520 nm in the IF plates. Uninfected primary cell cultures, as well as sera from SA-MCF-negative rabbits, were used as negative controls.

### 2.6 Immunohistochemistry

Immunohistochemistry (IHC) was performed on deer liver to determine the presence of inclusion bodies immunoreactive to an in-house hyperimmune serum made in rabbits inoculated with the OvHV-2 isolate. Deer liver sections were prepared on slides treated with 0.1% poly-l lysine (Sigmae Aldrich, St. Louis, Missouri, USA), deparaffinized, hydrated in alcohol baths, and subjected to antigen retrieval by incubation in citrate buffer (pH 6.0) in a pressure cooker for 9 min (24). Endogenous peroxidase was blocked with distilled water. water and H2O2 (3%) for 15 min. Non-specific proteins were blocked with powdered milk (4%). Primary incubation (24 h at 4 C) was with rabbit hyperimmune serum (1 in 5 dilution). Secondary incubation was with peroxidized anti-rabbit at a 1 in 50 dilution, later development was done with diaminobenzidine, counterstained with hematoxylin, and the slide was assembled. As a negative control, the primary antibody was not used in the liver and only the secondary antibody was added.

### 2.7 Nested endpoint polymerase chain reaction

The nested endpoint PCR was specific for the ORF75 segment of OvHV-2, which encodes the phosphoribosyl-formyl-glycinamidine synthase (FGARAT) that participates in purine metabolism and the production of viral tegument proteins (25).

The genetic material was extracted following the manufacturer’s instructions for the High Pure PCR kit of Roche® Campo Roche Diagnostics G (26). Primer pairs were designed by Baxter et al. (1993), and nested PCR was performed according to the method by Li et al. (1995), with some modifications (27,28). In the first amplification, primers SA-MCF 556 (5’ GTC TGG GGT ATA TGA ATCCAG ATG GCT CTC 3’) and SA-MCF 755 (5’ AAG ATA AGC ACC AGTTAT GCA TCT GAT AAA 3’) were used to amplify a 422-bp segment. The reaction conditions were adjusted to 25 μL, and a PCR Master Mix 5X (Taq-Load) was used. For the second amplification, the primers SA-MCF 555 (5’ TTC TGG GGT AGTGGC GAG CGA AGG CTT C 3’) and SA-MCF 556 (5’ GTC TGG GGT ATA TGA ATCCAG ATG GCT CTC 3’) were used to amplify a 238-bp segment (27). The concentration of external and internal primers was 10 μM. All reactions were performed in a GeneAmp® PCR System 9700 (Applied Biosystems, Waltham, MA, USA). Parameters for the PCR amplification were as follows: predenaturing at 95°C for 10 min, 35 cycles of denaturing at 94°C for 35 s, one alignment at 55°C for 1 min, one extension at 72°C for 45 s, and a final extension at 72°C for 10 min (14). In the second amplification, 1 μL of the first reaction was taken as a template for the second PCR round with the same parameters as those used for the first amplification round except for the alignment temperature, which was 60°C for 35 s.

We used 5 μL of the amplified product and 2 μL of the loading buffer with a red gel to visualize the reaction. The amplification product was subjected to electrophoresis for 35 min at 120 V in a 1.5% agarose gel. A positive result generated a 238-bp band in the second amplification.

A positive result generated a 238 bp band in the second amplification. A bovine blood sample positive by PCR and partial sequencing for OvHV-2 (ref. 6191), donated by the United States-Mexico Commission for the Prevention of Foot-and-Mouth Disease and Other Exotic Animal Diseases (CPA), was used as a positive control, and nuclease-free water was used as a negative control.

### 2.8 Partial sequencing of the ORF75 gene

For partial sequencing, the fourth blind passage of each of the six isolates obtained by inoculating the primary cell cultures with buffy coats from the different animal species affected by SA-MCF and the positive control donated by the CPA were used. In these primary cell cultures, 90 to 100% of the cell monolayer was destroyed 72 hours after infection. Samples were amplified with nested PCR to obtain a 422-bp fragment corresponding to a segment of the ORF75 gene (25). The amplification products were observed through electrophoresis in agarose gel, and the 422-bp band was visualized using UV transillumination. After visualizing the expected-size band, the amplification products were purified with the QIAquick® kit and quantified with a NanoDrop (Thermo Fisher Scientific) spectrophotometer. A total of 300 ng of each amplification was used, and external PCR primers were added. Sequencing was performed through capillary electrophoresis in a bidirectional manner with the Sanger sequencing method. The electropherograms of each pair of sequencing reactions were edited and assembled using the Biological Sequence Alignment Editor (BioEdit), v 7.2.0; (29) and sequences were aligned using MEGA 11 software (30).

### 2.9 Phylogenetic analysis

The seven consensus sequences obtained with BioEdit software (29) were compared with those reported in GenBank, selecting those with an identity percentage greater than 95%, using BLAST (Basic Local Alignment Search Tool) from the NCBI (National Center for Biotechnology) (31). The obtained sequences were phylogenetically analyzed with sequences reported in GenBank for the Macavirus genus (OvHV-2, AIHV-1, and AIHV-2) and human gammaherpesvirus types 8 and 4 (HVH-8 and HHV-4) using the maximum likelihood method with 1000 bootstrap repetitions. The evolutionary distances were calculated with the 3-parameter Tamura method, and the analysis was carried out with Mega 11 software (30).

## 3. Results

### 3.1 Clinical disease and pathology

In September 2018, an outbreak of SA-MCF occurred in the CEIEPAA, Tequisquiapan, Queretaro, Mexico. The affected animals showed mucogingival ulcers in the mouth vestibule and tongue, hypersalivation, corneal opacity, lower food consumption, and weight loss of variable severity (Table 1). In addition, fourteen deer died during the SA-MCF outbreak due to the severity of the ulcers in the oral mucosa that hindered them from eating (Figure 2).

**Figure 2.**
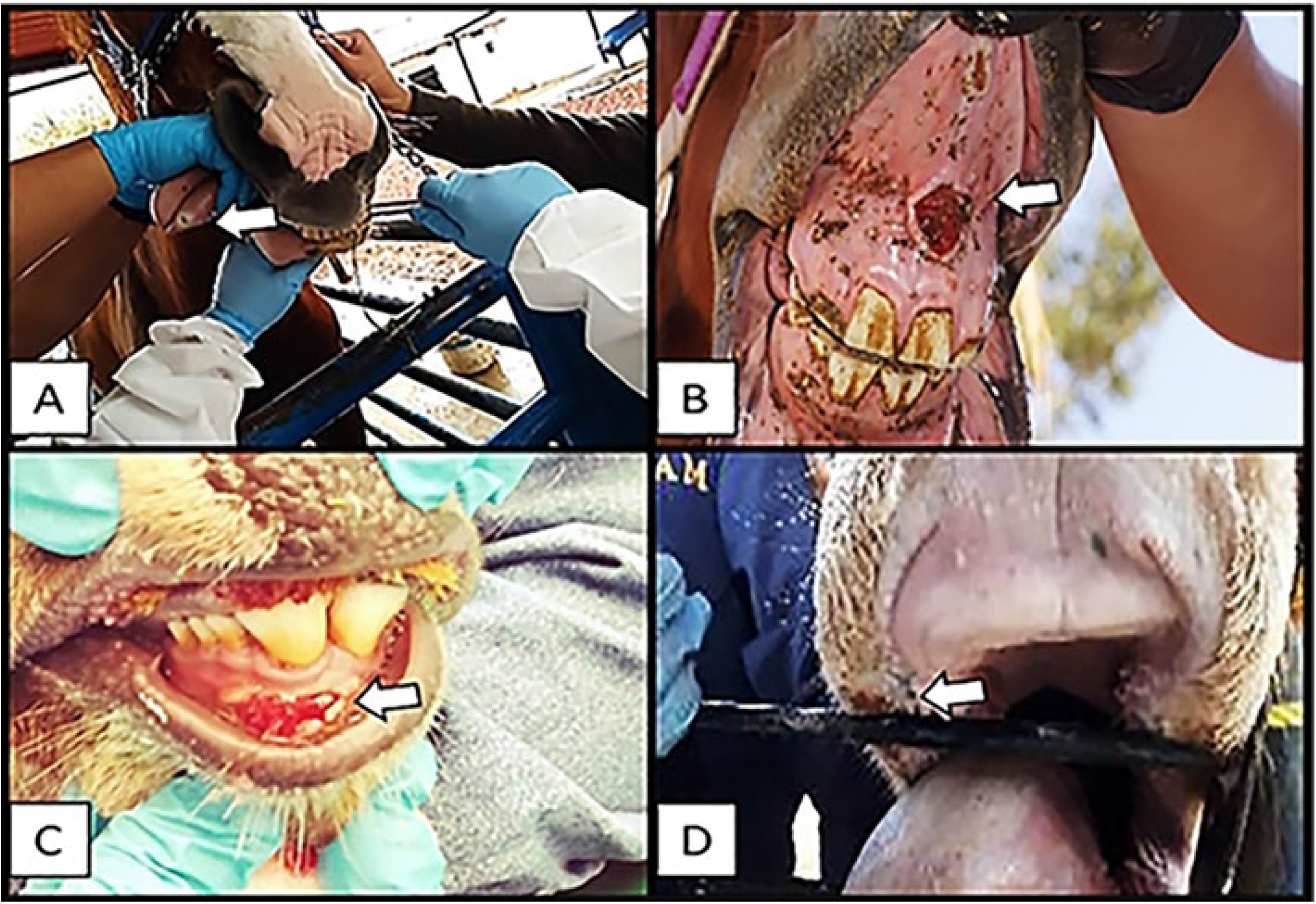
Oral mucosa lesions observed in the animals suffering from MFC, showing erosions and ulcers, as well as bleeding of the oral mucosa and base of the tongue (white arrows). A) and B) horses, C) deer and D) dairy cow.

Of the 14 deer that died during the outbreak, the one with the highest number of clinical signs and lesions was chosen, and the corresponding postmortem examination revealed marked lymphadenomegaly of retromandibular lymph nodes with congested and edematous, dorsally ulcerated tongue, focal ulcers on the rumen pillars covered by a pseudomembrane, and slightly enlarged and congested liver. The primary microscopic lesions observed were vasculitis in the tongue and perivasculitis in the rumen and tongue, corneal edema and corneal neovascularization, retromandibular lymph nodes with lymphoid hyperplasia, cytoplasmic eosinophilic inclusion bodies in the liver, tongue with thrombosis and lymphoplasmacytic and lymphoblastic perivascular infiltrates, and rumen with perivascular mononuclear infiltrate mainly composed of lymphocytes, plasma cells, and lymphoblasts (Figure 3). No postmortem lesions were observed in the lung.

**Figure 3.**
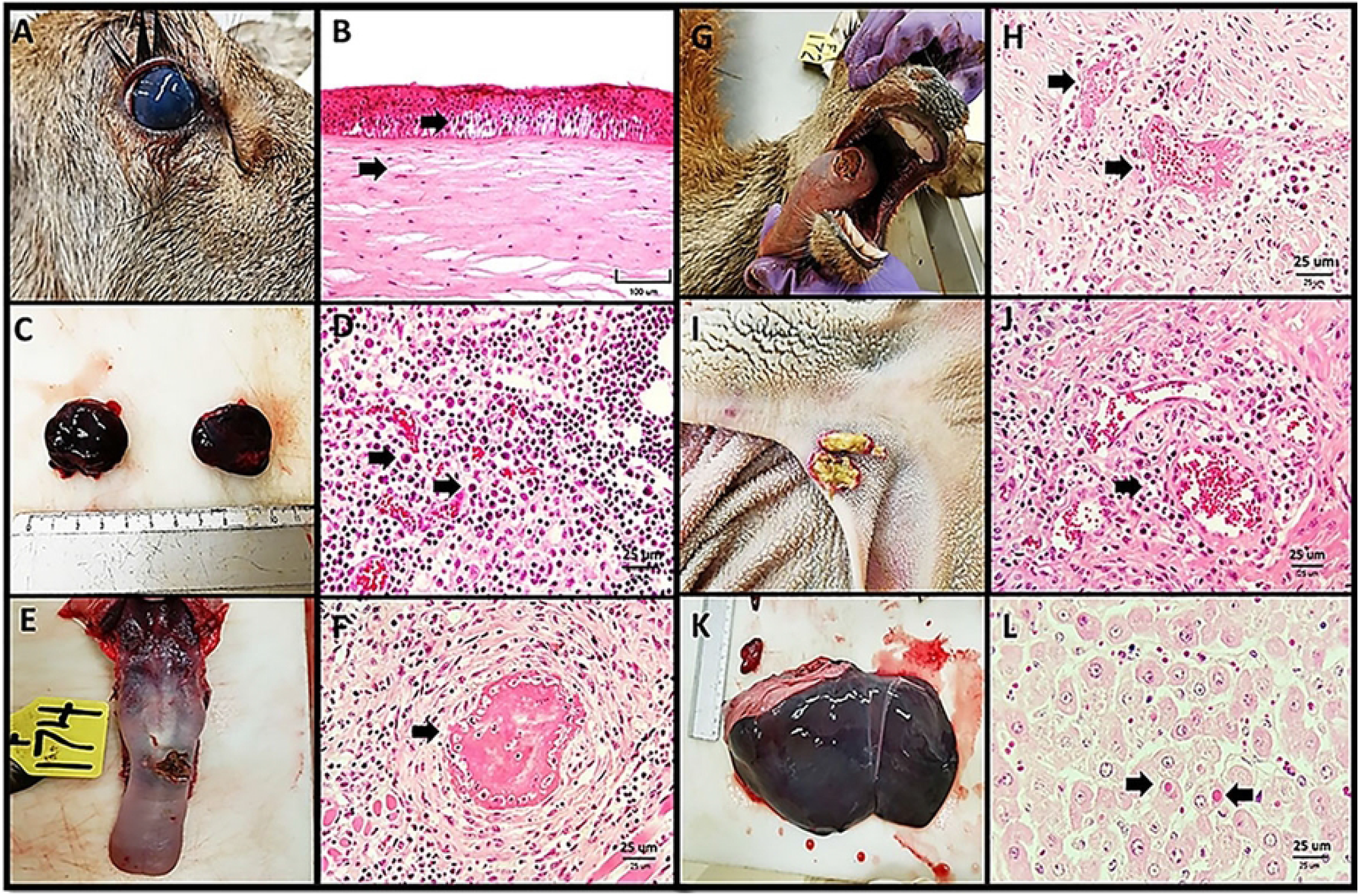
Postmortem lesions observed in affected deer in the SA-MCF outbreak in Queretaro, Mexico. A- Deer with corneal opacity, B- Cornea with edema and detachment of the corneal epithelium, C- Enlarged, congested and edematous retromandibular lymph nodes, D- Parafollicular region with numerous lymphocytes, plasma cells and lymphoblasts, E- Tongue showing loss of the continuity of the epithelium (ulcer) in the dorsal portion, F-Tongue that presents a blood vessel with a fibrin thrombus that totally occludes the lumen and fibrinoid necrosis, G-Tongue with an ulcer in the dorsal portion of the tongue, with food encrustation, H- Tongue with thrombosis and lymphoplasmacytic and lymphoblastic perivascular infiltrate, I-Rumen with focal ulcers in the pillars of the rumen, covered by a pseudomembrane, J-Rumen with perivascular inflammation composed of lymphocytes, plasma cells and lymphoblasts, K-Congested liver and enlarged (hepatomegaly), L- Liver with areas of necrosis and cytoplasmic eosinophilic inclusion bodies.

### 3.2 OvHV-2 detection in a buffy coat by PCR

A total of 213 buffy coat samples from the animals with clinical signs compatible with SA-MCF were tested by nested end-point PCR; 86 were positive (40.38%), and 127 were negative (59.62%). The PCR results revealed that the resident goats and sheep had the highest percentage of animals positive for OvHV-2, whereas the dairy cattle and horses had the lowest number of positive animals (Table 1).

### 3.3 Virus isolation in primary cell cultures from the buffy coat

To isolate OvHV-2, six pools of endpoint PCR-positive buffy coat samples from each affected species were used (deer, dairy cattle, feedlot cattle, sheep, goats, and horses) (Table 1). PCR-positive buffy coat samples from resident goats and goats in the quarantine area were mixed in a single pool. Each buffy coat group pool from the different animals was processed separately, and four blind passages were performed.

The cytopathic effect (CPE) caused by the buffy coat pools was similar in all six isolates and was characterized for generating small clusters of refractile cytomegalic cells; this CPE was observed between 48 and 72 hours postinfection (Figure 4).

**Figure 4.**
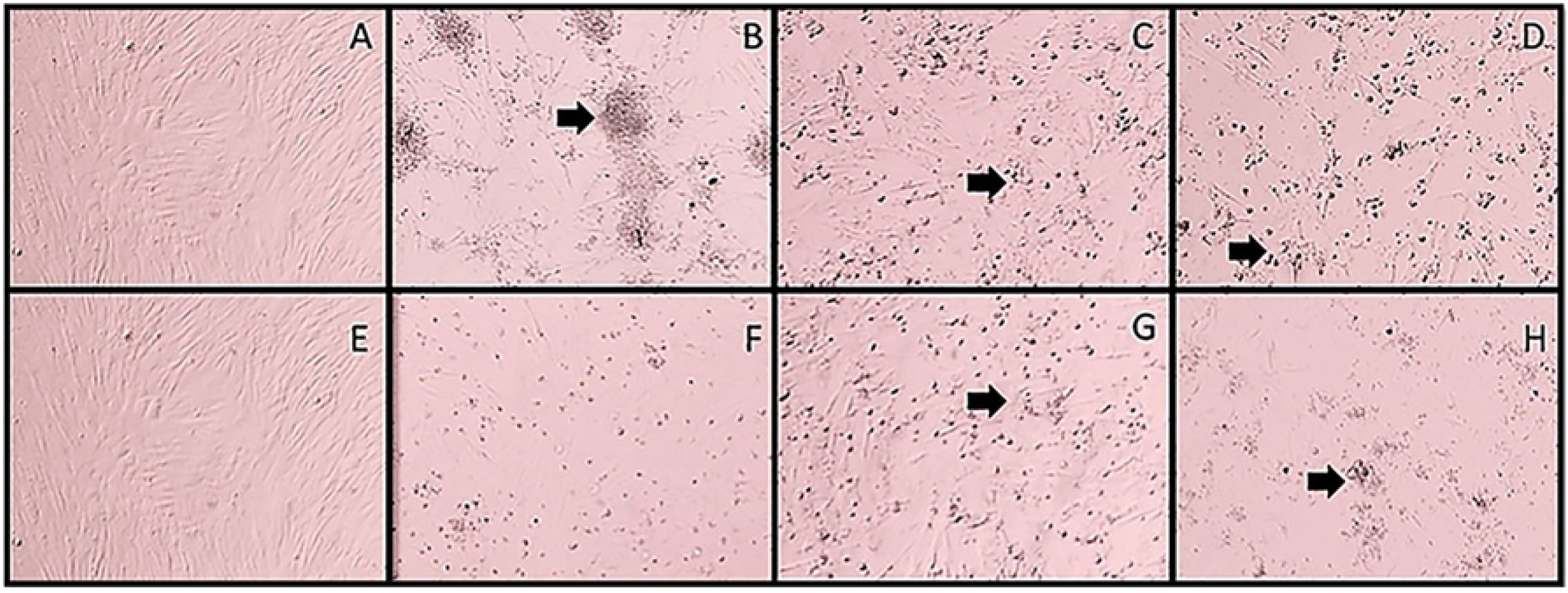
Cytopathic effect in primary cell cultures of rabbit testicle cells infected with buffy coats of the different animal species affected during the SA-MCF outbreak at 72 hours postinoculation. A and E) Negative controls, B) horse isolation, C) sheep isolation, D) goat isolation, F) deer isolation, G) dairy cattle isolation, H) feedlot cattle isolation. Arrows indicate small clusters of refractile cytomegalic cells. Magnification: 10X.

The isolate from sheep and goats showed marked CPE with 50% destruction of the monolayer from the first passage to 72 hours postinfection and complete destruction of the monolayer between 96 and 120 hours postinfection. Likewise, the percentage of monolayer CPE increased from the second pass and fourth pass onward for the goat isolate and the sheep isolate, respectively (Table 2). On the other hand, primary cell cultures infected with buffy coat pools of horses, dairy cattle, feedlot cattle, and deer showed little CPE, with monolayer destruction between 20 and 30% in the first and second passages, respectively. In contrast, in the third and fourth passages, destruction of the monolayer between 50 and 90% was observed at 72 hours postinfection (Table 2).

**Table 2.**
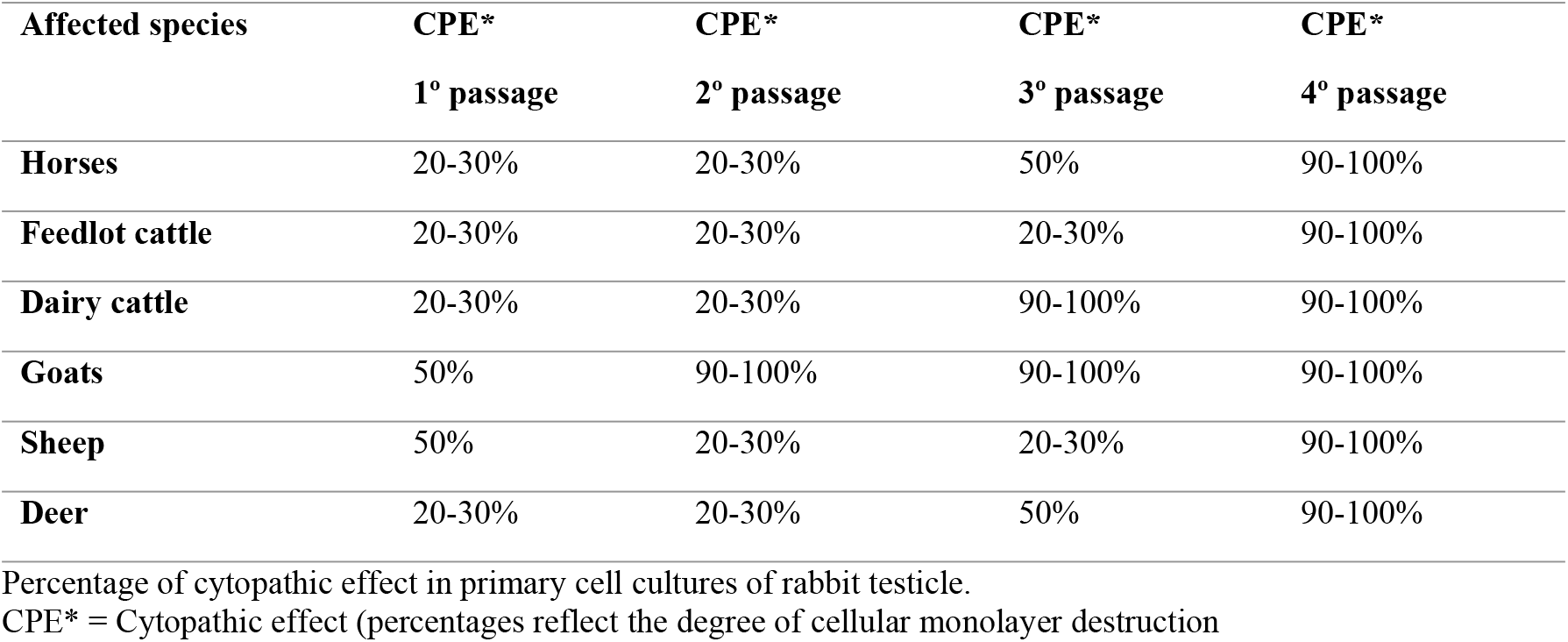
Cytopathic effect in primary cell cultures.

| Affected species | CPE* | CPE* | CPE* | CPE* |
| --- | --- | --- | --- | --- |
|  | 1° passage | 2° passage | 3° passage | 4° passage |
| **Horses** | 20-30% | 20-30% | 50% | 90-100% |
| **Feedlot cattle** | 20-30% | 20-30% | 20-30% | 90-100% |
| **Dairy cattle** | 20-30% | 20-30% | 90-100% | 90-100% |
| **Goats** | 50% | 90-100% | 90-100% | 90-100% |
| **Sheep** | 50% | 20-30% | 20-30% | 90-100% |
| **Deer** | 20-30% | 20-30% | 50% | 90-100% |

Percentage of cytopathic effect in primary cell cultures of rabbit testicle.
CPE\* = Cytopathic effect (percentages reflect the degree of cellular monolayer destruction)

The isolate obtained from buffy coats of the goats was the best adapted to the primary cell cultures of the rabbit testicle, showing a marked CPE starting at the first passage and increasing along the following blind passages. Viral titers of the isolated viruses from horse, goat, sheep, deer, feedlot cattle and dairy cow on cell cultures varied between log 10^3^ TCID_50_%/mL and log 10^4.19^ TCID_50_%/mL.

### 3.4 Inclusion bodies

Primary cell cultures were infected with four blind passages of the six isolates; 24 h post-infection, the plates were fixed with 4% paraformaldehyde and H&E stained to observe the inclusion bodies. In each infected well, cytoplasmic acidophilic inclusion bodies enveloped in a white halo were observed; in most, the nucleus was displaced (Figure 5). In the wells infected with the four blind passage of the isolate from goats and horses, more than ten cytoplasmic acidophilic inclusion bodies were observed; in contrast, in the wells infected with the isolates from dairy cattle, beef cattle, deer and sheep, only four or five acidophilic inclusion bodies could be observed throughout the well.

**Figure 5.**
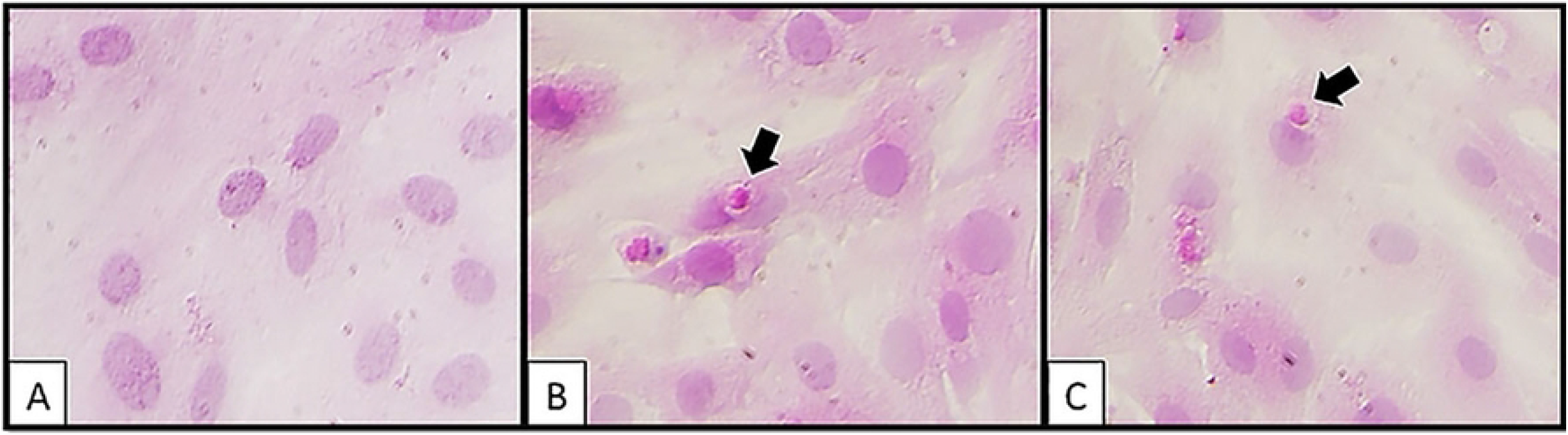
Cytoplasmic acidophilic inclusion bodies were observed at 24 h postinfection and stained with hematoxylin/eosin. A) Negative control, B) Second passage of the horses’ isolate, C) Second passage of the goats’ isolate. The black arrows indicate the inclusion body surrounded by a white halo located in the perinuclear area. Magnification: 20X.

### 3.6 Immunofluorescence and immunoperoxidase

Cells from the primary cell cultures of rabbit testes infected with the third passage of the six isolations showed specific marks by IF techniques using the hyperimmune polyclonal serum against OvHV-2. The IF technique evidenced antigen-antibody binding as an apple-green emission at 520 nm and by the IP technique as a brown precipitate. Isolates from horses and goats presented the highest proportion of cells with intracytoplasmic immunoreactivity, between 40% and 50% of the monolayer, respectively (Figure 6).

**Figure 6.**
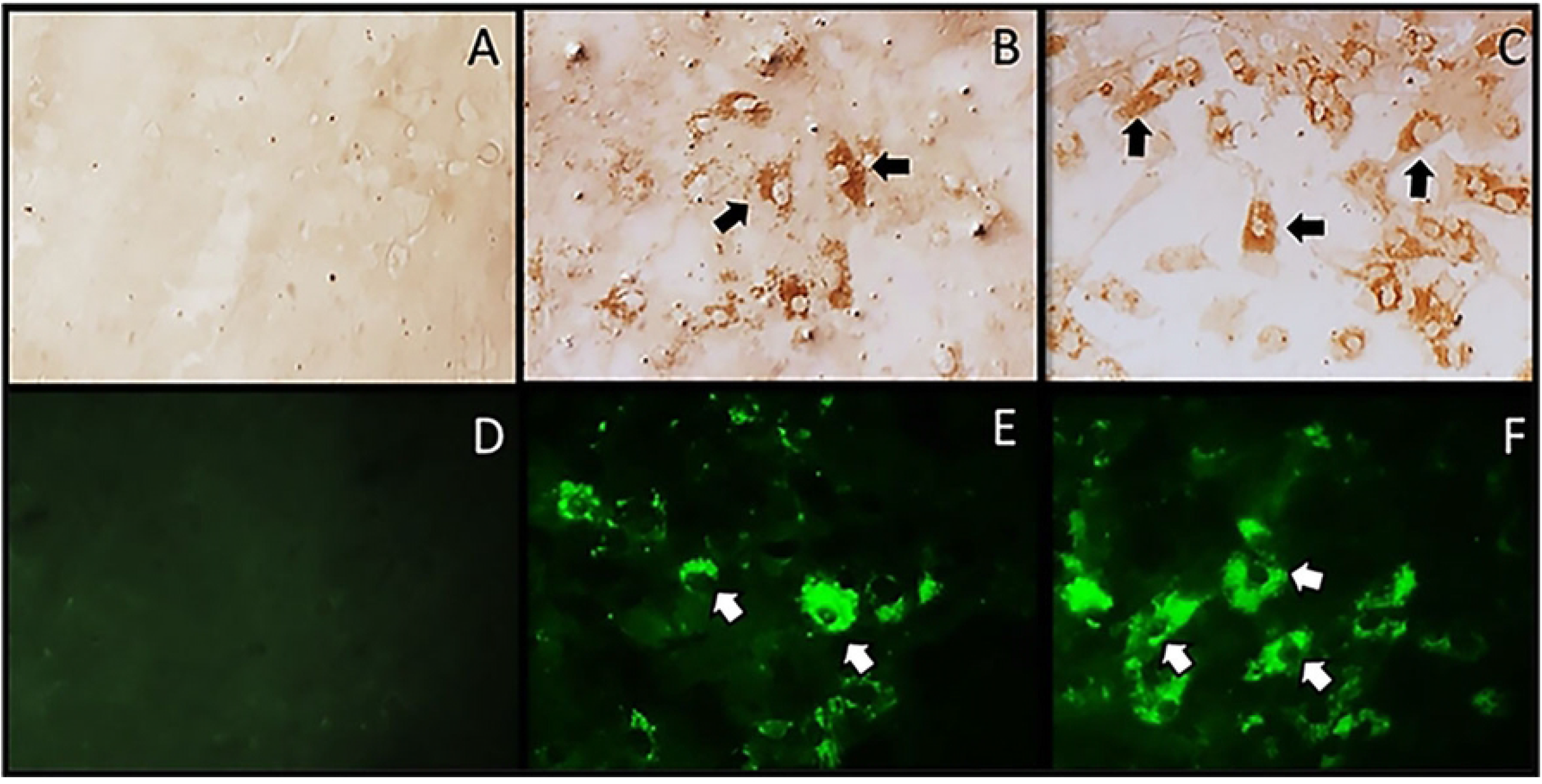
Cytoplasmic immunoreactivity with the IF and IP techniques in primary cell cultures of rabbit testicle cells. A) IP negative control, B) IP horse isolate (third passage), C) IP goat isolate (third passage); D) IF negative control, E) IF horse isolate (third passage), F) IF goat isolate (third passage). Arrows indicate the cytoplasmic immunoreactivity evidenced as an insoluble brown color for the IP and a green-apple color for the IF. Magnification: 20X.

### 3.7 Deer liver immunohistochemistry

Positive immunoreactivity to anti-OvHV-2 hyperimmune rabbit serum was observed in the cytoplasmic inclusion bodies of hepatocytes from affected deer during the MCF outbreak. In addition, positive intracytoplasmic and slightly intranuclear immunoreactivity was observed in hepatocytes (Figure 7).

**Figure 7.**
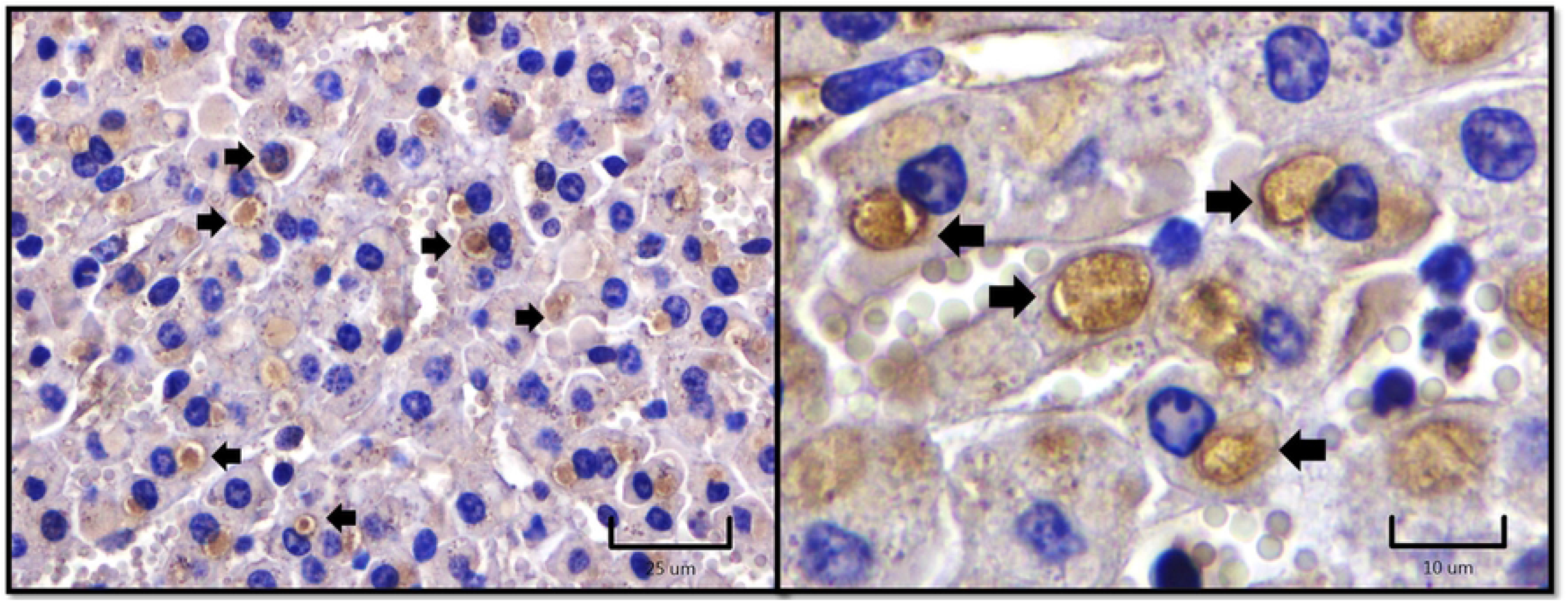
Deer liver immunohistochemistry. Immunoreactivity to OvHV-2 antigens with rabbit hyperimmune serum, in cytoplasmic inclusion bodies (black arrows), within the cytoplasm and in the nucleus of hepatocytes (yellow arrows).

### 3.8 Nested end-point PCR on OvHV-2 isolates

DNA extraction was performed from the supernatants of the four cell passages of the six isolates obtained from the buffy coats of the different species affected during the SA-MCF outbreak. The nested end-point PCR revealed a 422-bp band in the first amplification and one of 238 bp in the second amplification. Figures 8 and 9 show the 238 bp bands corresponding to isolates positive for OvHV-2 by the nested endpoint PCR technique.

**Figure 8.**
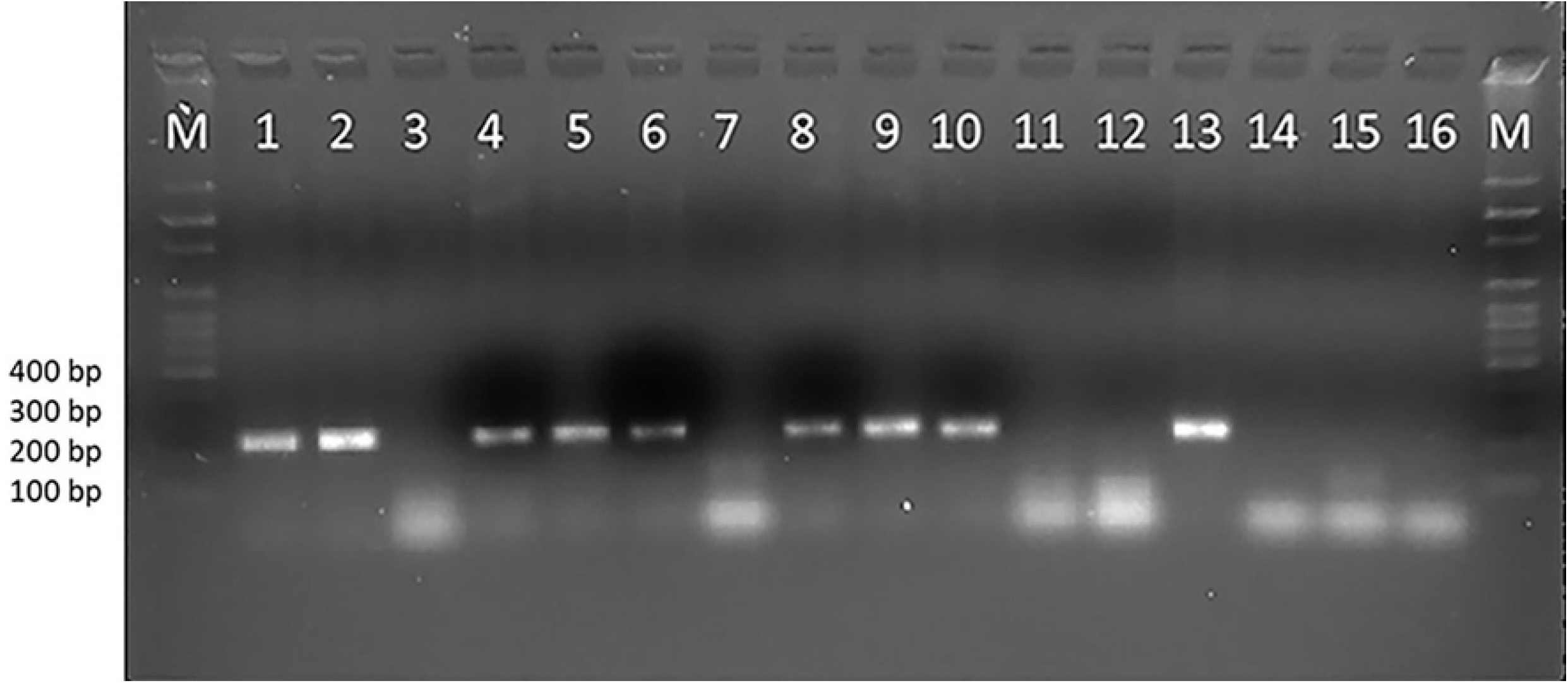
Agarose gel (1.5%) run in TAE 1X buffer. Nested PCR of the first and second passages of primary cell cultures of rabbit testicles. Lane M- Marker of molecular weight of 100 bp. Lane 1- Horses’ isolation passage 1. Lane 2- Horses’ isolation passage 2. Lane 3- Sheep isolation passage 1. Lane 4- Sheep’ isolation passage 2. Lane 5- Goat isolation passage 1. Lane 6- Goat isolation passage 2. Lane 7- Deer’ isolation passage 1. Lane 8- Deer’ isolation passage 2. Lane 9- Dairy cattle isolation passage 1. Lane 10-Dairy cattle isolation passage 2. Lane 11- Feedlot cattle isolation passage 1. Lane 12- Feedlot cattle’s isolation passage 2. Lane 13- Positive control. Lane 14- Negative control of amplification (CNA). Lane 15- Negative control of extraction (CNE). Lane 16- Noninfected primary cell cultures (CNCP). Lane M- molecular weight marker.

**Figure 9.**
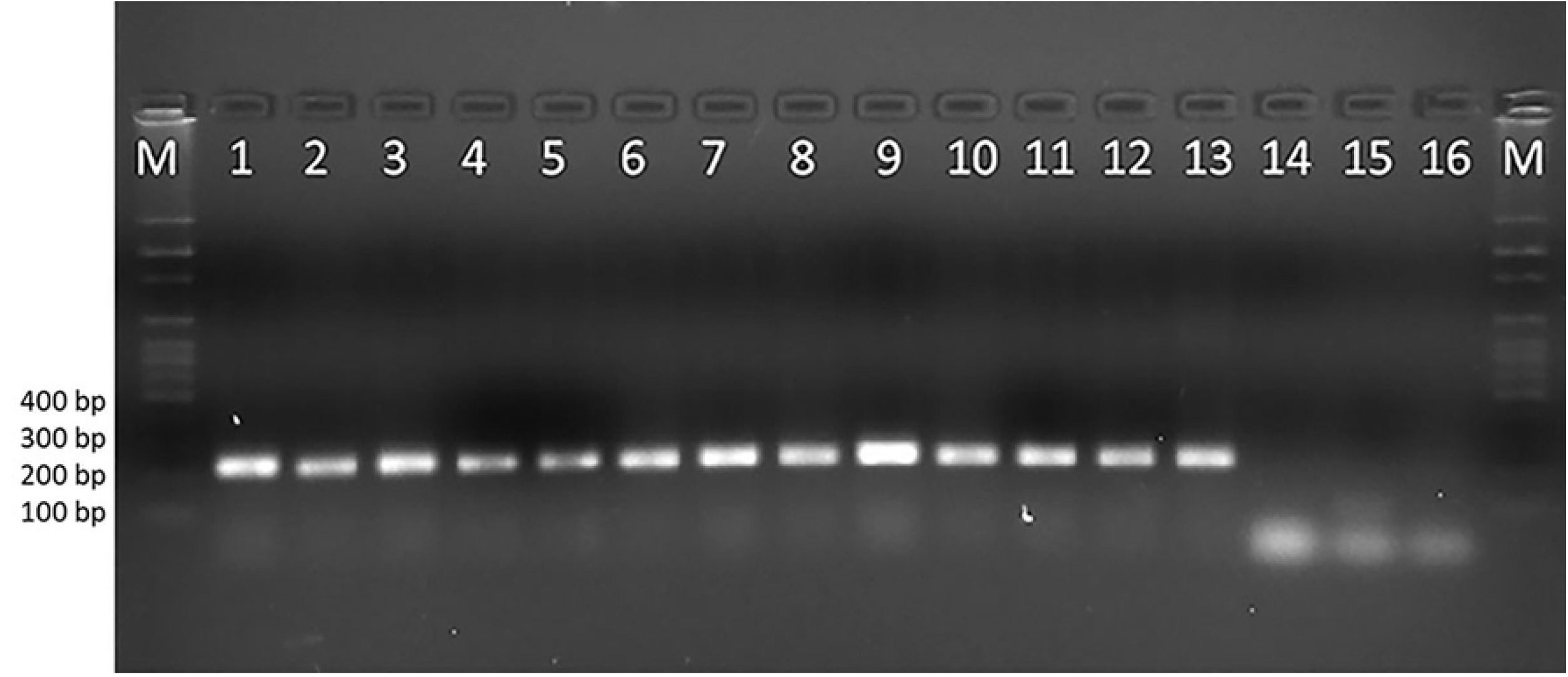
Agarose gel (1.5%) run in TAE 1X buffer. Nested PCR of passages three and four in primary cell cultures of rabbit testicles. Lane M- Marker of molecular weight of 100 bp. Lane 1- Horses’ isolation passage 3. Lane 2- Horses’ isolation passage 4. Lane 3- Sheep isolation passage 3. Lane 4- Sheep isolation passage 4. Lane 5- Goat isolation passage 3. Lane 6- Goat isolation passage 4. Lane 7- Deer’ isolation passage 3. Lane 8- Deer’ isolation passage 4. Lane 9- Dairy Cattle isolation passage 2. Lane 10- Dairy Cattle isolation passage 4. Lane 11- Feedlot cattle isolation passage 3. Lane 12- Feedlot cattle isolation passage 4. Lane 13- Positive control. Lane 14- Negative control of amplification (CNA). Lane 15- Negative control of extraction (CNE). Lane 16- Noninfected primary cell cultures (CNCP). Lane M- molecular weight marker.

Isolates from horses, goats, and dairy cattle were positive from the first passage onward; isolates from sheep and deer were positive from the second passage onward (Figure 8).

However, from the third and fourth passages, all six isolates were positive by nested PCR and generated a 238 bp band during the second amplification (Figure 9).

### 3.9 Phylogenetic analysis

Alignment of the 422-bp partial sequences of the ORF75 gene indicated that the seven sequences obtained from the six isolates and the positive control shared 98 and 99% nucleotide identity with OvHV-2, as well as with the sequences obtained from the GenBank database. Regarding the structure of the inferred topology, the analysis indicated that our seven partial sequences constituted a clade with different OvHV-2 ORF75 genes reported in other regions of the world and separate from the *Alcelaphine gammaherpesvirus* and *Human gammaherpesvirus*, which were in different clades (Figure 10).

**Figure 10.**
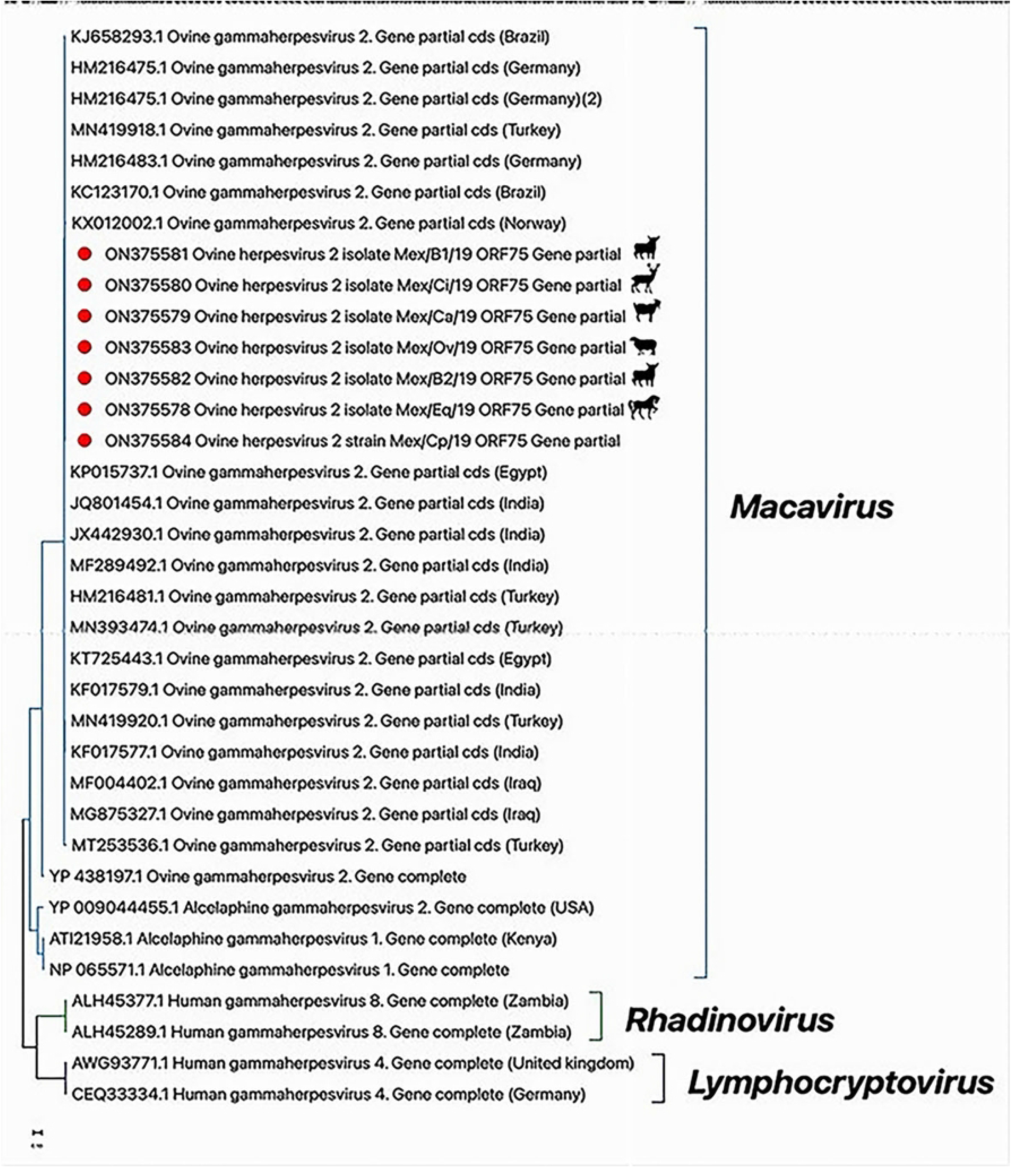
Phylogenetic analysis of the partial sequences of the ORF75 gene. The tree is at scale, and the bar indicates the number of substitutions per site. The Mexican isolates are marked with red rhombus.

## 4. Discussion

In this study, we have described an outbreak of SA-MCF that occurred in the CEIEPAA, located in Tequisquiapan, state of Querétaro, Mexico. In addition, we isolated and identified OvHV-2 by different diagnostic techniques, including immunofluorescence, immunoperoxidase, PCR end point and partial sequencing of the ORF75 gene. During the outbreak, many animals of different species were affected, showing typical head and eye clinical signs compatible with SA-MCF and corneal opacity (8). SA-MCF was confirmed using a PCR method to detect OvHV-2.

In our study, we used buffy coats from affected animals to isolate OvHV-2, which is similar to the study by Hristov et al. (2016), who used bison buffy coats and whole blood, lung, and spleen samples from different affected animals during an outbreak of SA-MCF (18). From the third passage in the primary cell cultures of rabbit testis, a marked cytopathic effect was observed at 72 hours post-infection consisting of small foci of refractile cytomegalic cells. This CPE was also described by Rossiter et al. (1980) during the isolation of AIHV-1 and by Hristov et al. (2016) during the isolation of OvHV-2 from camels affected by SA-MCF (18,32).

In the present study, titers of viruses isolated from horses, goats, sheep, deer, beef cattle, and dairy cattle in primary cell cultures of rabbit testes ranged from log 10^3^ and log 10^4.19^ TCID_50_%/mL; this titer differs from the one found by Hristov et al. (2016), who described a titer of 10^6^ TCID_50%_/mL in Madin-Darby bovine kidney (MDBK) after passage 19 (18). Based on the titer they found, our low titer could be due to its predominant cell association in the primary isolation in cell cultures, and after five passages, there is a period during which the viral genome undergoes rearrangements, leading to attenuation and resulting in better yields of cell-free virus (33).

We identified perinuclear and cytoplasmic acidophilic inclusion bodies by H&E and immunohistochemistry, in the liver of deer that died during the SA-MCF outbreak and in primary cell cultures from rabbit testes infected with OvHV-2 isolates.

These results coincide with those found by Aluja et al. (1969), who described cytoplasmic acidophilic inclusion bodies in neurons of cows infected with SA-MCF (21). Likewise, our results are similar to those reported by Goss et al. (1947), who described these inclusion bodies in Purkinje cells and epithelial cells of mucous membranes (34). These results differ from those of Hristov et al. (2016), who observed only intranuclear inclusion bodies in infected cell lines (18). The presence of perinuclear and cytoplasmic acidophilic inclusion bodies observed in the present study could be due to sites of accumulation and retention of overproduced viral and cellular proteins; these macromolecular structures called “dense bodies” are very common in human cells infected by cytomegalovirus. Most of these structures are sequestered in Golgi-derived vesicles, and some dense bodies are located near or in contact with nuclear membranes, where they appear to inject viral proteins necessary for viral assembly through nuclear pores (35). The accumulation of viral protein in the cytoplasm also coincided with the description by Nelson et al. (2013), who showed that the distribution of the capsid protein is cytoplasmic in OvHV-2 infected terminal hosts, suggesting that there is abortive viral replication (6). Therefore, we can speculate that cytoplasmic inclusion bodies suggest an accumulation of viral proteins necessary for virion assembly, in addition to abortive virions and cellular proteins. In any case, more studies are needed to confirm this hypothesis.

Importantly, we observed intracytoplasmic immunoreactivity through IF and IP in the primary cell cultures; these results agree with those reported by Headley et al. (2022), who identified intracytoplasmic immunoreactivity in multiple tissues affected by OvHV-2 using a monoclonal antibody (mAb-15A) targeted at a conserved epitope among all Macaviruses (36). Finally, OvHV-2 was identified in the primary cell cultures through endpoint nested PCR targeting a 238 bp fragment of the ORF75 gene, with a greater intensity starting in the third and fourth passages in all isolates from the different affected species. These results indicate that to generate many free viruses and achieve adaptability of the virus to the cell, several blind passes are needed, coinciding with the report by Hristov et al. (2016) (18)

The definitive diagnosis confirmation of OvHV-2 infection was performed using partial sequencing of the 422-bp fragment of the ORF75 gene. With this approach, it was possible to determine that all obtained isolates had 98 to 99% nucleotide identity among them and with the OvHV-2 sequences reported in different regions of the world. These results agree with those reported by Martins et al. (2017), who sequenced 13 positive samples of bovines from different areas of Brazil and found 97 to 100% similarity with the OvHv-2 of several other regions of the world (37).

In conclusion, the SA-MCF outbreak in Tequisquiapan, Queretaro, Mexico, was caused by OvHV-2, and the disease was identified by clinical signs, microscopic lesions, viral isolation, inclusion bodies, IF, IP, IHC, PCR, and partial sequencing of the ORF75 gene. Furthermore, this study demonstrated for the second time that horses are susceptible to OvHV-2 infection and can develop clinical disease. Consequently, SA-MCF should be considered in the differential diagnosis of vesicular diseases of this species. Finally, it was determined that primary cell cultures from rabbit testes are susceptible and permissible to viral replication.

## Funding information

This study was financed by PAPIIT-UNAM (IN213321), PAPIIT-UNAM (IT201918) and T.L. Madrigal-Valencia received a scholarship SEP-CONACYT, ID (968751).

## Acknowledgments

Thanks to the Special Cathedra “Dr. José E. Mota” 2019-2020. The authors thank the United States-Mexico Commission for the Prevention of Foot-and-Mouth Disease and Other Exotic Animal Diseases (CPA) for their technical support in laboratory testing and positive control.

## Notes

### Competing Interest Statement

The authors have declared no competing interest.

